# Silver nanoparticle biosynthesis utilizing *Ocimum kilimandscharicum* leaf extract and assessment of its antibacterial activity against certain chosen bacteria

**DOI:** 10.1101/2023.12.05.570271

**Authors:** Horyomba Siaka Ouandaogo, Souleymane Diallo, Eddy Odari, Johnson Kinyu

## Abstract

Using plants in the biological production of silver nanoparticles (AgNPs) is becoming increasingly important as a practical and environmentally benign method. In the current work, silver nanoparticles were made utilizing the significant *Ocimum kilimandscharicum*, and their potential to combat certain bacteria was discussed. Both aqueous and methanol plant extracts were used for reducing silver nitrate at different time intervals (30 to 150 minutes) and pH (2 to 11). The UV-visible absorption spectrum recorded for both methanol and aqueous extracts revealed successful synthesis of AgNPs. The antimicrobial activity of the AgNPs was evaluated against *Escherichia coli* ATCC 25922, *Salmonella choleraesuius* ATCC 10708, and *Staphylococcus aureus* ATCC 25923. 50mg/ml was the concentration of the extracts tested. The best zone of inhibition was recorded for the methanol and aqueous-mediated AgNPs, ranging from 12±1 to 16±1mm. The methanol and aqueous extract silver nanoparticles had the same Minimum Inhibitory Concentration (6.25±0.00 mg/ml), whereas the Minimum Bactericidal Concentrations were 12.5±0.00 and 25±0.00 mg/ml, respectively. The highest inhibition zone of 16±1 mm was observed against *Salmonella choleraesuius* with 50±0.00 mg/ml aqueous silver nanoparticles. The results show that the silver nanoparticles made with *Ocimum kilimandscharicum* have potent antibacterial action against those microorganisms.

## 1. Introduction

*Ocimum kilimandscharicum*, a member of the Lamiaceae family, is indigenous to East Africa and is grown elsewhere in the world. The taste of this species is strong but less appealing. It is a pubescent understory shrub with quadrangular branchlets and a fragrant smell. This plant, which may reach 2.44 m, is easily identified by its shrubby form [1]. It has ovate leaves and pale-yellow blooms [2]. This plant gained popularity since it is a source of camphor. This plant is frequently used in traditional medicine to cure a broad range of conditions, including diarrhoea, measles, stomach discomfort, and colds and coughs. The leaves have also been used as a therapy for measles and to alleviate congested chest, cough, and colds [2]. The plant extracts have been found to have wound-healing properties [3]. Plant extracts with biologically active components contain insecticidal, fungistatic [4], and antibacterial properties [5]. Crop protection, controlled pesticide release, target gene transfer, plant hormone administration, and using nanosensors for early disease detection in plants are just a few of the many applications of nanotechnology in agriculture [6–8]. It is being investigated that nanoparticles may be used as a secure and reliable management technique to control phytopathogens [9].

Due to their distinct physiochemical features, silver nanoparticles are well recognized for displaying high antimicrobial action against bacteria [10], fungi [11] and viruses [12]. A wide variety of human infections have been targeted through the extensive use of silver nanoparticles’ antimicrobial properties [13]. Many researchers have argued that silver nanoparticles are effective against a wide variety of plant infections because they are antibacterial and antifungal [14].

Metallic nanoparticle manufacturing via chemical and physical means is frequently expensive and involves potentially harmful substances. Numerous biological methods have been created as an alternative to create safe, affordable, and environmentally acceptable nanoparticles [15]. It is anticipated that employing plants for biological synthesis would offer several advantages over bacteria and fungi. The accessibility of plants and the existence of a wide variety of metabolites that help produce silver nanoparticles are two of these advantages.

The use of *O. kilimandscharicum* in the manufacture of AgNPs was not reported in the literature. The goal of the current work was to bio-synthesize AgNPs using *O. kilimandscharicum* and assess its efficacy in the management of bacterial infections.

## 2. Materials and methods

### 2.1. Chemicals

Legacy Lab Africa Ltd., the Kenyan company, provided all chemicals utilized in this investigation. Analytical-grade chemicals were employed throughout.

### 2.2. Preparation of plant extract

*Ocimum kilimandscharicum* leaves were collected from the Juja region in Kenya in March 2021 and transported to the Pan African University of Basic Science, Technology and Innovation, Kenya. Mr John Kamau Muchuku, Department of Botany, Jomo Kenyatta University of Agriculture and Technology, identified and verified the plant leaf material, and a voucher specimen (HSO – JKUATBH/001/A – 2021) was put in the herbarium for reference.

Fresh leaves were surface cleaned with running water and rinsed with distilled water. The leaves were then chopped into little pieces, allowed to air dry at room temperature in the shade, and ground into a fine powder with an electric blender. Following extraction with 2000 ml of methanol (70%), 100 g of the finely powdered leaf extract was placed in an orbital shaker incubator at 25°C for 72 hours. A rotatory evaporator was used to concentrate the methanolic extract filtrate. 100 g of the finely powdered leaves extract was placed in 2000 ml distilled water at 80°C. After 1 hour, the mixture was filtrated. Whatman filter paper number 1 was used to filter the extract and the filtrate [16]. The resultant extracts were stored in a refrigerator at 4 °C.

### 2.3. Green synthesis of silver nanoparticles

#### 2.3.1. Preparation of the 1 mM AgNO_3_ solution

We utilized the approach of Ghabban *et al.* [17] with a few minor adjustments. Briefly, 1.7 g of silver nitrate was dissolved in 1000 ml of deionized water in a 1000 ml volumetric flask and then transferred into an amber bottle to create a stock solution of 10 mM of AgNO3 aqueous solution.

From the stock, 1 mM of AgNO3 aqueous working solution was created by measuring 10 ml of 10 mM AgNO3 and adding 90 ml of distilled water. The flask was wrapped in aluminium foil to avoid photoreduction and kept in the dark.

#### 2.3.2. Synthesis of silver nanoparticles

Three millilitres of plant extract (methanolic and aqueous) and one hundred millilitres of silver nitrate solution were combined in a conical flask (1 mM). The bioreduction (Ag+→Ag0) process was allowed to occur by covering the conical flask with aluminium foil and incubating it at room temperature [18]. Visual changes in colour and absorbance readings taken with a UV-vis spectrophotometer covering the range of 200 nm to 800 nm confirmed the presence of silver nanoparticles (AgNPs) in the reaction mixture. The solution was centrifuged at 18,500 rpm for 60 minutes at 4 °C to remove impurities and freeze-dried. Two rounds of centrifugation were used to achieve purification [18].

### 2.4. Synthesized silver nanoparticles characterization

#### 2.4.1. Analysis of UV-Visible spectroscopy

PAUSTI’s UV-Visible spectrophotometer (Model 6800, Jenway) was used to evaluate the reduction of silver nitrate ions (Ag+) by *Ocimum kilimandscharicum* leaf extract at wavelengths between 800 and 300 nm [19,20]. According to Singh *et al*. [13], and Shaik *et al.* [21] phenolics, flavonoids, and tannins play a vital role during redox reaction and act as capping agents. The 400-500 nm peak, a typical peak of silver nanoparticles owing to surface resonance plasmon (SRP), was used to validate the synthesis of silver nanoparticles.

#### 2.4.2. Analysis of functional group using FTIR

Fourier infrared spectrometry was used to classify functional groupings (FTIR, Shimadzu 8400, Kyoto, Japan). We combined 10 mg of powdered AgNPs with 100 mg of Potassium Bromide to make the salt disc. A range of 4000 to 500 cm-1 was used to capture the spectra [20].

#### 2.4.3. Dynamic Light Scattering

Malvern ZETASIZER NANO was utilized to determine the particle size, polydispersity index (PDI), as well as stability of biosynthesized AgNPs (Model: ZEN5600, Serial number: MAL1168679) [22].

### 2.5. Antimicrobial screening of the extracts and their silver nanoparticles

#### 2.5.1. Organisms testing

*O. kilimandscharicum* leaf extracts and their silver nanoparticles were examined for their antibacterial activity against Gram-positive and negative bacteria strains. All strains are American Type Culture Collection (ATCC). The Kenya Medical Research Institute (KEMRI) donated the bacterial strains, *E. coli (ATCC 25922)*, *S. choleraesuius (ATCC 10708*), and *S. aureus (ATCC 25923*).

#### 2.5.2. Bacterial cultures, upkeeping and preparation of the inoculum

On nutrient agar, the chosen bacteria have been subcultured and maintained. The bacterial strains were cultured overnight in Muller Hinton broth (MHB) (0.5 McFarland standard).

#### 2.5.3. Zone Of Inhibition (ZOI)

The Agar diffusion technique was used to test the antibacterial activity of the crude extract and silver nanoparticles displayed by Ali *et al.*, Ahmad *et al.*, Shanmugapriya *et al.* [23–25] with some modifications. 5.6 grams of Muller Hinton Agar was dissolved in 200 millilitres of distilled water, then heated and autoclaved (parameters). The media (20 ml) was placed into each petri dish and allowed to solidify, and newly subcultured bacteria using sterile swabs, were transferred to the plate. Six (6) mm wells were drilled into each plate using a sterile cork borer. In brief, 100 µl of the prepared extract solution (50 mg/ml) and (50 mg/ml) of silver nanoparticles were pipetted into the wells and allowed to diffuse. As the standard, 10 mg of gentamycin was utilized as a positive control. After incubating the plates for 24 hours, the inhibition zones were evaluated (mm).

#### 2.5.4. Determination of the Minimum Inhibitory Concentration (MIC)

Using the resazurin-based 96-well plate microdilution method, the minimum inhibitory concentration of *O. kilimandscharicum* extract and its mediated silver nanoparticles were measured in triplicates [26]. Various doses (200 mg/ml to 0.39 mg/ml) were generated using two-fold dilution. 200 µl of newly cultivated bacteria were pipetted into each well, followed by 200 µl of the test samples. The plates were incubated for 24 hours at 37 °C. The next day, 100 µl of resazurin dye was pipetted into each well and incubated for an additional two hours. The minimum inhibitory concentrations were determined by observing the colour changes of each well (from blue to pink) to identify the lowest inhibitory concentrations [27].

#### 2.5.5. Determination of the Minimum Bactericidal Concentration (MBC)

The least inhibitory concentration is the lowest concentration at which colony growth is inhibited. Extracts of *O. kilimandscharicum* and silver nanoparticles mediated were tested to find their bactericidal concentrations using the approach given by Lemma *et al.* and Mkangara *et al.* [28,29] with some modifications. Using a sterile swap, the well contents (3 wells downstream) in which no colour changes had occurred were moved to fresh sterile Petri plates and infected. It was cultured for a further twenty-four hours and examined for bacterial growth. The least bactericidal concentration from the 96 plates was extrapolated to the plates without observable growth.

### 2.6. Statistical analysis

The Mean ± standard error (SE) was used to express the experimental results. R studio, Excel, and SAS9.2 were used to perform an analysis of variance (ANOVA). The significance difference between the means was performed using Turkey test. P value less than 0.05 were considered significant.

## 3. Results and discussion

### 3.1. Silver synthesis and characterization

There are several ways to synthesize nanoparticles, including chemical, physical, and biological approaches. Although physical and chemical techniques are the most popular, they are both damaging to the environment and costly [30]. Plants are abundantly accessible, inexpensive, and contain bioactive substances [31]; hence, green nanoparticle manufacturing has lately attracted considerable attention. Moreover, using plants to synthesize is safe and ecologically beneficial [13,32]. Plants are rich in secondary metabolites which usually facilitate the synthesis of silver nanoparticles. In a research study conducted by Saxena *et al.* [33], flavonoids, alkaloids, glycosides, saponins, and tannins were present in methanolic extract of *O. kilimandscharicum*.

#### 3.1.1. UV-Visible spectroscopy

An aliquot of the solution was analyzed using UV-visible spectrophotometry to track the development of silver nitrate bioreduction. Figures 2 and 3 distinctive display peaks measured at 424 nm for the methanolic extract and 442 nm for the aqueous extract [34,35], respectively.

**Figure 1.**
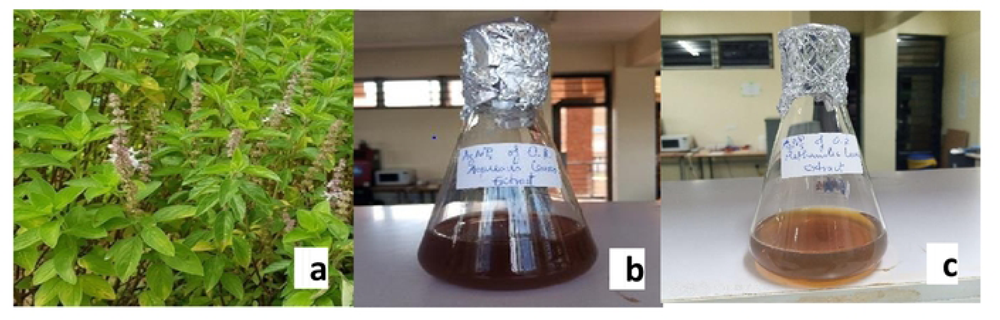
a, *O. kilimandscharicum* plant; **b,** synthesized AgNPs using the aqueous extract; **c,** synthesized AgNPs using the methanolic extract.

**Figure 2.**
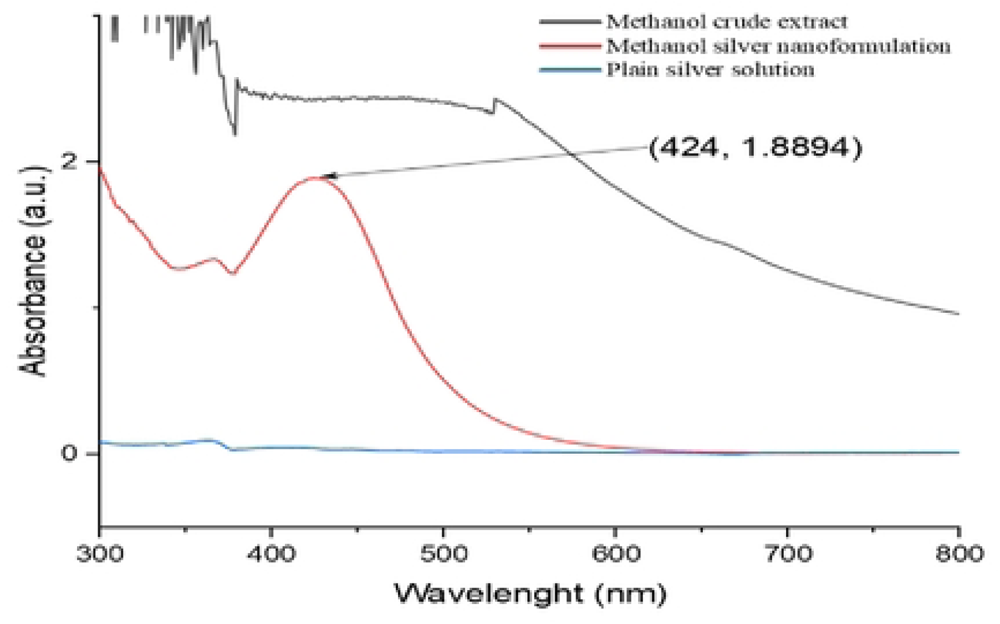
UV/Vis spectra of *O. kilimandscharicum* methanolic extract-mediated AgNPs

**Figure 3.**
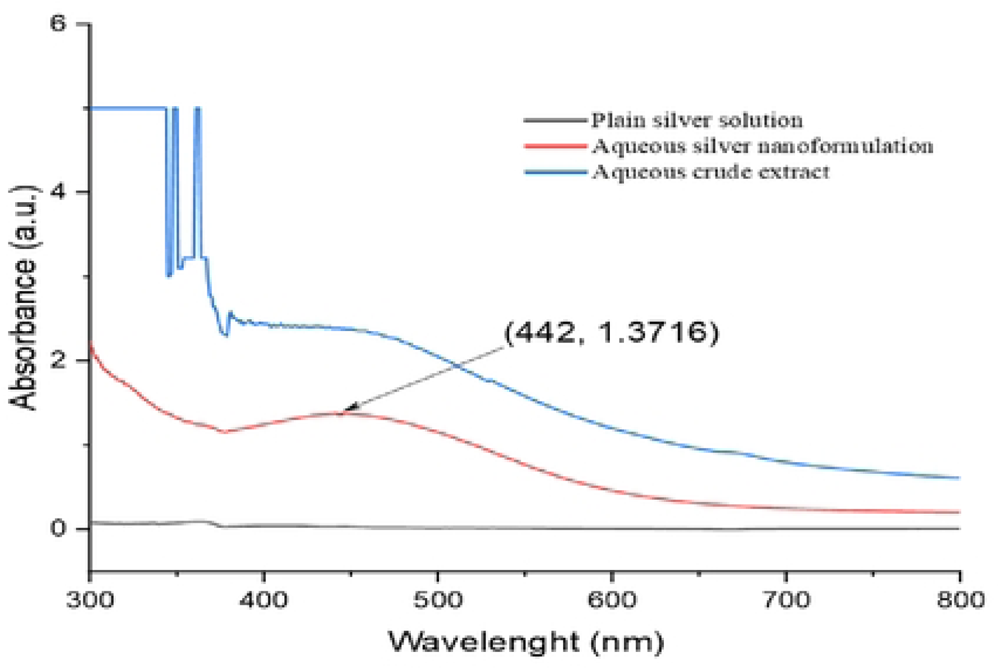
UV/Vis spectra of *O. kilimandscharicum* aqueous extract-mediated AgNPs

##### 3.1.1.1. Produced AgNP’s stability over time

AgNPs suspension was subjected to different time conditions and measured the absorbance spectra of UV/Vis in a scan range of 350nm to 650nm. The times tested were 30mn, 60mn, 90mn, 120mn, and 150mn (Figures 4 and 5).

**Figure 4.**
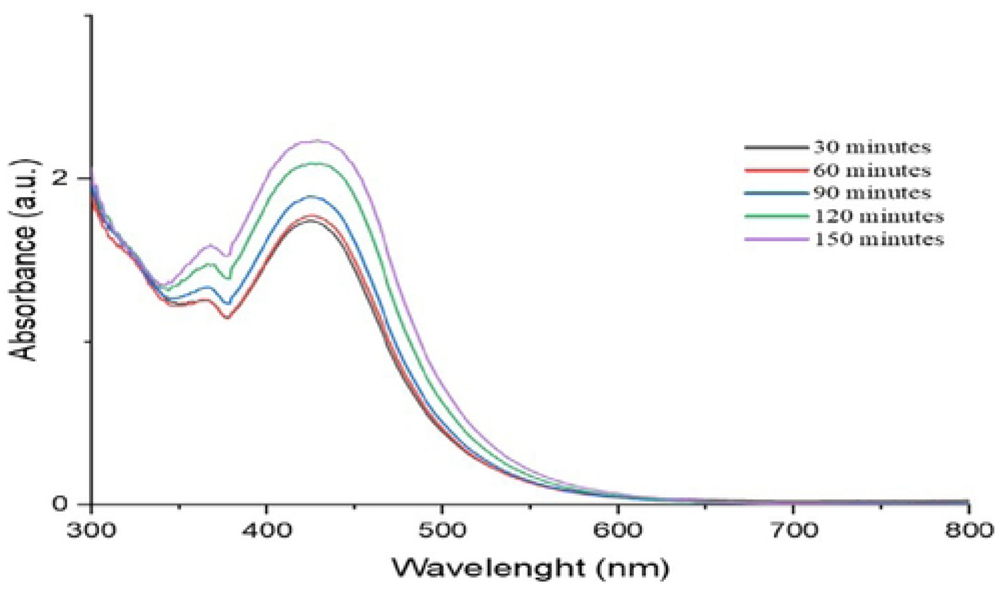
UV/Vis spectra of time stability of O. *kilimandscharicum* methanolic extract-mediated AgNPs

**Figure 5.**
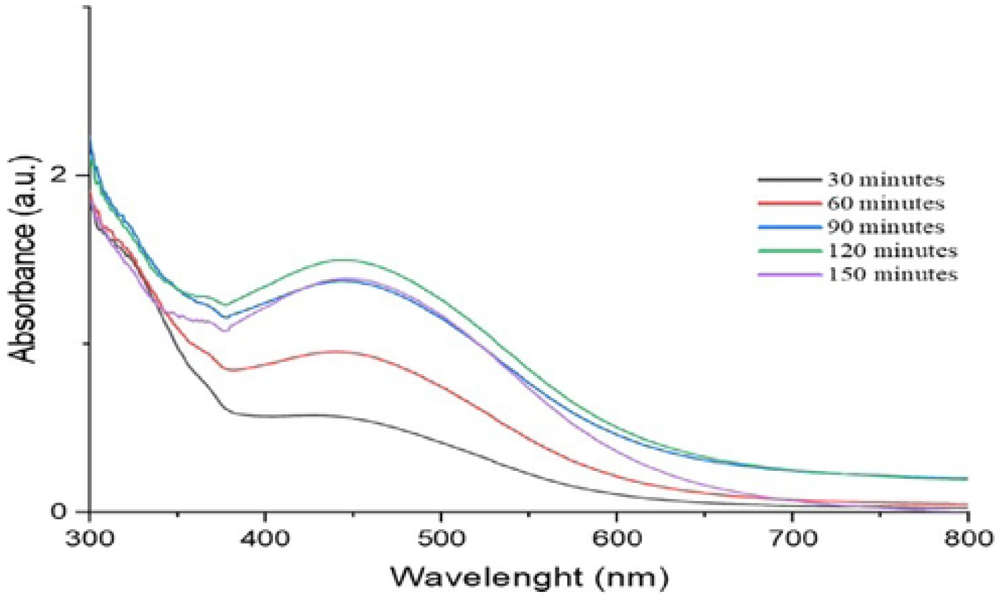
UV/Vis spectra of time stability of O. *kilimandscharicum* aqueous extract-mediated AgNPs

Ninety minutes (90 min) is the best time to stop our synthesis process for the AgNPs using *O. kilimandscharicum* methanolic leaves extract, corresponding to a sharp peak (Figure 4). In comparison, 120 minutes is the best time for *O. kilimandscharicum* aqueous extract-mediated AgNPs (Figure 5).

##### 3.1.1.2. Produced AgNP’s stability pH

The produced AgNPs suspension was aliquoted into 5 test tubes, each holding about 3 ml of the AgNPs suspension. The suspensions in the test tubes were adjusted to various pH settings ranging from 2 to 11 using drops of either 1N NaOH or 1N HCl. The absorbance was measured from 300 to 800 nm (Figures 6 and 7).

**Figure 6.**
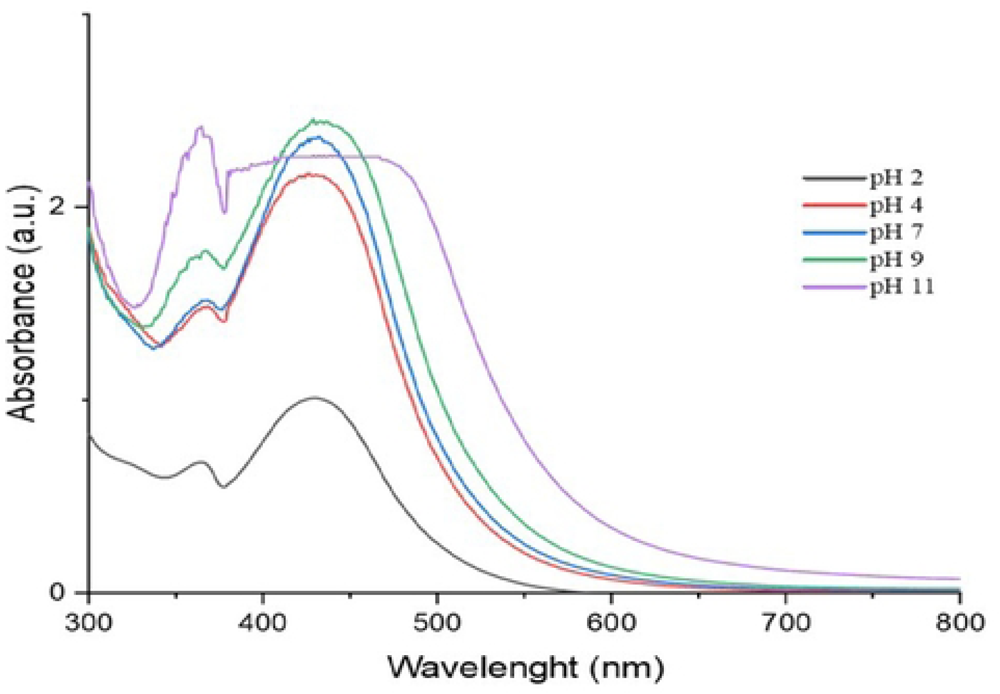
UV/Vis spectra of Ph stability of O. *kilimandscharicum* methanolic extract-mediated AgNPs

**Figure 7.** UV/Vis spectra of Ph stability of *O. kilimandscharicum* aqueous extract-mediated AgNPs

Both AgNPs synthesized using the methanolic and aqueous extracts are good at pH 2, 4, and 7, showing that the AgNPs are stable at these pHs (Figures 6 and 7). Above pH 7, the AgNPs are no longer stable, as revealed by the very broad peaks (Figures 6 and 7).

Our findings are consistent with those of Ndikau *et al.* [36] and Khan *et al.* [37], which saw maxima at 433 nm and 424 nm, respectively. The surface of silver nanoparticles often exhibits recognizable peaks between 400 nm and 500 nm due to Plasmon resonance excitation [30].

#### 3.1.2. Analysis of functional groups using FTIR

Using Fourier Transform Using infrared analysis, the functional groups involved in reducing and capping silver nanoparticles were identified. Based on the FT-IR measurements, both main and minor peaks were determined.

Possibly owing to O-H or N-H stretching vibrations of polyphenols, flavonoids, and alkaloids, the peaks at 3348.19 cm-1 moved to 3325.05 cm-1, and 3355.91 cm-1 shifted to 3348.19 cm-1 in the AgNPs (Figures 8 and 9) (Table 1). Khanal *et al.* obtained a similar outcome [38]. Initial phytochemical data supported the findings because polyphenols are known to be natural reducing agents [39].

**Figure 8.**
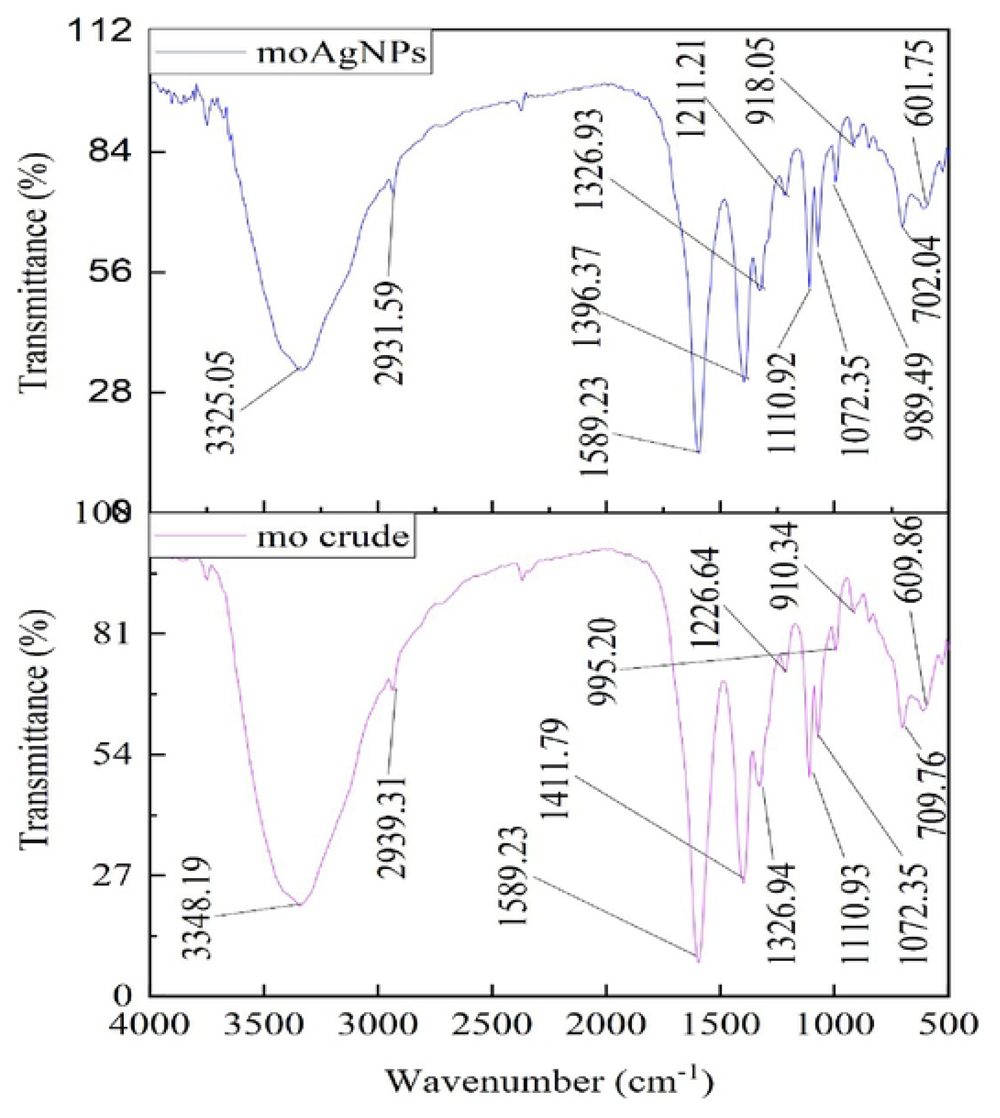
FT-IR spectra for the methanolic extract of *O. kilimandscharicum* and AgNPs

**Figure 9.**
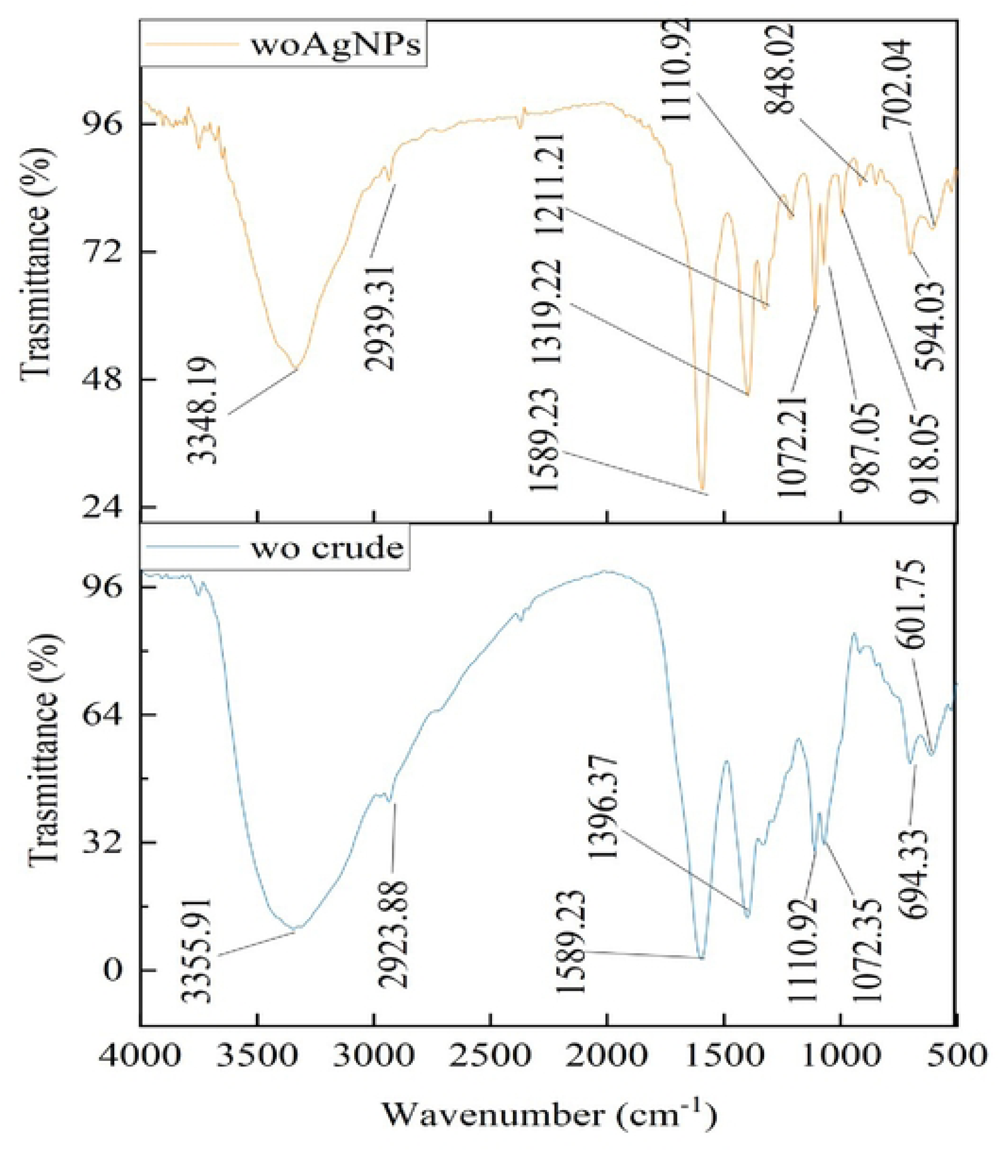
FT-IR spectra for the aqueous extract of *O. kilimandscharicum* and AgNPs

**Table 1.** FTIR analysis of methanol and aqueous extract and its mediated silver nanoparticles.

| Frequency(cm<sup>1</sup>) | Chemical bond | Assignment | Extracts | AgNPs |
| --- | --- | --- | --- | --- |
| 3500–3200(s,b) | O–H stretch, N–H stretch | Alcohols, phenols<br>flavonoids/ alkaloids | 3348.19,<br>3355,91 | 3325.05,<br>3348.19 |
| 3300-2500(m) | O-H stretch | Carboxylic acids | 2923.28,<br>2939.31 | 2939.31,<br>2931,59 |
| 1650–1585(m) | N–H bend | 1 amines | 1589.23,<br>1589.23 | 1589.23,<br>1589.23 |
| 1500-1400(m) | C=C stretch | Aromatics | 1411,79 |  |
| 1335–1250(s) | C–N stretch | Aromatic amines | 1326.94 | 1319.22,<br>1326.93 |
| 1320–1000(s) | C–H stretch | Alcohols, carboxylic acids, esters, ethers | 1110.92,<br>1072.35,<br>1226.64,<br>1110.93,<br>1072.35 | 1319.22,<br>1211.21,<br>1110.92,<br>1072.21,<br>1072.35 |
| 1000–650(s) | =C-H bend | Alkenes | 910.34, 694.33,<br>709.76 | 987.05,<br>918.05,<br>848.02,<br>702.04 |
| 640–515(m) | =C-Br | Alkyl halides | 609.86, 601.75 | 594.03,<br>601.75 |

As a result of carboxylic acids, the spectral bands between 3300 and 2500 cm-1 have been attributed to O-H stretching. Similar to the findings published by Ahmad *et al.* [24], we observed a peak at 2939.31 cm-1, 2931,59 cm-1, and 2923.28 cm-1 (Figures 8 and 9) (Table 1).

The band 1589.23 cm1 was detected between 1650 and 1585 cm-1 (Figures 8 and 9) (Table 1), and this may be related to the N-H bending of 1° amine. Comparatively, the findings of these investigations were comparable to those published by Ahmad *et al.* [24], Hanna *et al.* [40], and other researchers, Pungle *et al.* [18]. C=C stretching vibrations attributed the peak at 1411.79 cm-1 to the aromatic compounds.

The detected band at 1110.92 cm -1 is due to the C-N stretch of an aliphatic amine group in the protein [41]. Simultaneously, many spectral bands, 1110.92, 1226.64, 1110.93, 1072.35, 1319.22, 1211.21, 1072.21 cm-1, resulting from C-H stretch owing to alcohols, carboxylic acids, esters, and ethers, as they lie between 1320 and 1000 cm-1, were detected.

Due to alkene and C–Br stretching due to alkyl halides, C-H bending corresponds to spectral bands at 987.05, 918.05, 848.02, 702.04, 594.03, and 601.75 cm -1, respectively. As shown by FTIR analysis, polyphenols, alkaloids, flavonoids, and proteins have a crucial role in decreasing, capping, and stabilizing biosynthesized AgNPs. Proteins contribute to the reduction, stabilization, and prevention of nanoparticle aggregation [42,43]. This reinforces the stability, high surface charge, and superb colloidal nature of dynamic light scattering.

#### 3.1.3. Dynamic Light Scattering

Figures 10a and 10b depict the results for the average particle size and stability of produced AgNPs from the methanolic extract, while Figures 11a and 11b show the average particle size and stability of produced AgNPs from the aqueous extract, respectively.

**Figure 10a.**
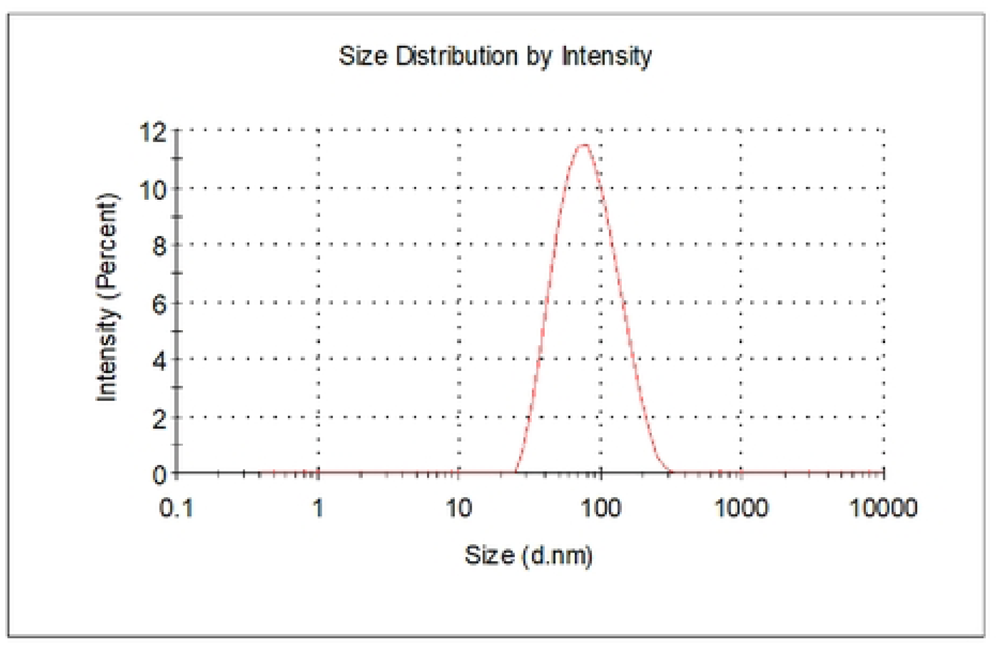
size distribution of AgNPs of *Ocimum kilimandscharicum* methanolic leaf extracts

**Figure 10b.**
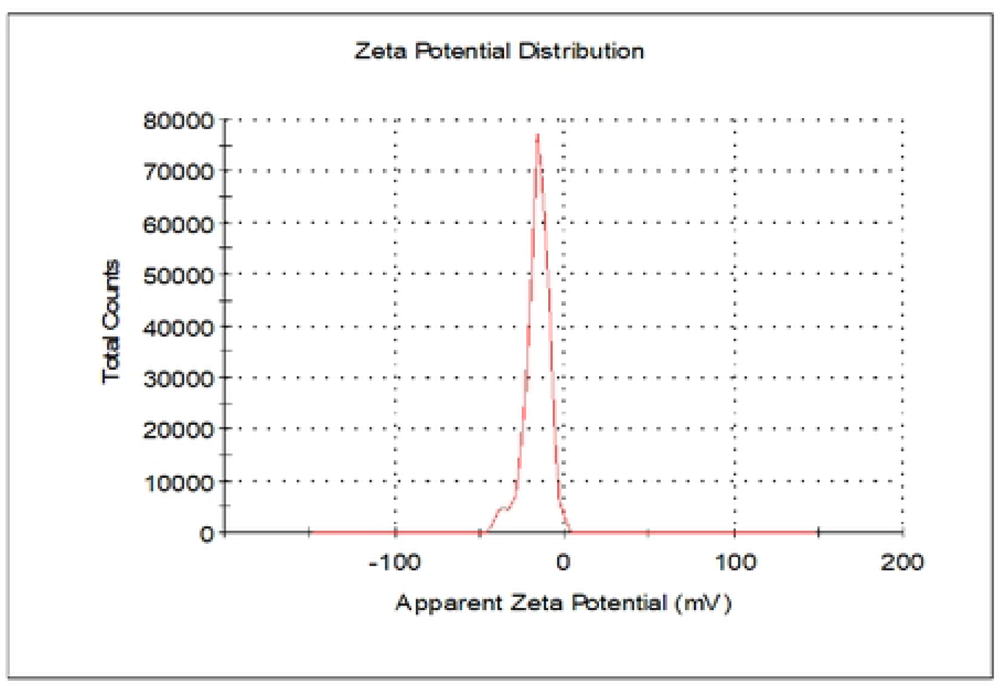
zeta potential distribution of AgNPs of *Ocimum kilimandscharicum* methanolic leaf extracts

**Figure 11a.**
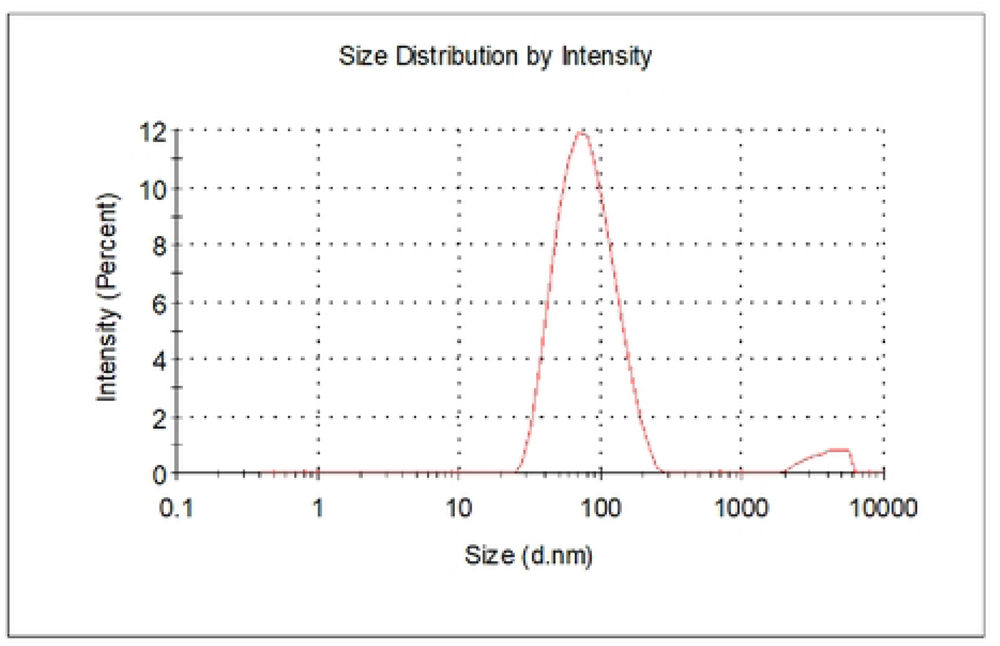
size distribution of AgNPs of *Ocimum kilimandscharicum* aqueous leaf extract

**Figure 11b.**
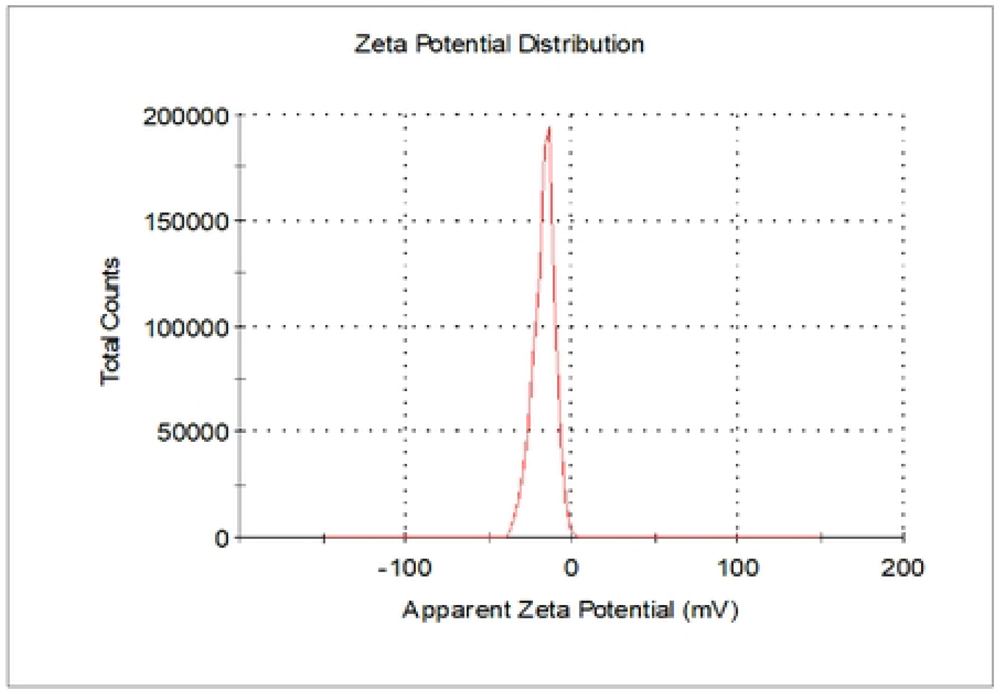
zeta potential distribution of AgNPs of *Ocimum kilimandscharicum* aqueous leaf extracts

Nanoparticle sizes are typically between 1 and 100 nm [40]. The nanoparticles’ positivity and negativity determine their aggregative quality. When two charges coexist, they tend to attract one another, resulting in their accumulation [44]. The *O. kilimandscharicum*-mediated AgNPs from the methanolic extract had an average particle size of 69.72±0.45 nm, a polydispersity index of 0.27±0.026, and a zeta potential of -17.6±1.89 mV. For the *O. kilimandscharicum*-mediated AgNPs from the aqueous extract, the average particle size was 70.14±1.15 nm, the polydispersity index was 0.27±0.005, and the zeta potential was -18.1±2.47 mV as determined by dynamic light scattering (Table 2, respectively). Negative charges suggested that the biosynthesized nanoparticles lacked clustering and clumping characteristics [45]. Studies have demonstrated how a positive charge of + 5.68 mV resulted in agglomerative solid potential [46]. Similar results were also reported; -24.6 mV [47], -30 mV [48], -22.3 mV [49], and -22.7 mV [45]. The polydispersity index (PDI) for monodispersed particles ranges from 0.01 to 0.70, while polydispersed particles have more than seven (7) polydispersity index values [45]. Our findings indicated nanoparticles to be promising in terms of particle size, stability, and dispersion.

**Table 2.** Characterization of the silver nano formulated from methanolic and aqueous extracts of *Ocimum kilimandscharicum* leaves by zeta.

| Solvent<br>extract | Size distribution (d. nm)<br>±SE | Polydispersity index<br>(PDI) ±SE | Zeta potential distribution<br>(mV) ±SE |
| --- | --- | --- | --- |
| Methanol | $69.72 \pm 0.45$ | $0.27 \pm 0.026$ | $-17.6 \pm 1.95$ |
| Water | $70.14 \pm 1.15$ | $0.27 \pm 0.005$ | $-18.1 \pm 2.47$ |

### 3.2. Antimicrobial screening of the extracts and their silver nanoparticles

#### 3.2.1. Zone Of Inhibition (ZOI)

In the present study, we evaluated the antibacterial activity of silver nanoparticles synthesized from the methanolic and aqueous leaf extracts of *O. kilimandscharicum* and its crude extract against Gram-negative (*Escherichia coli ATCC 25922, Salmonella choleraesuius ATCC 10708*) and Gram-positive bacteria (*Staphylococcus aureus ATCC 25923*). Figures 12, 13, and Table 3 show the findings, respectively.

**Figure 12.**
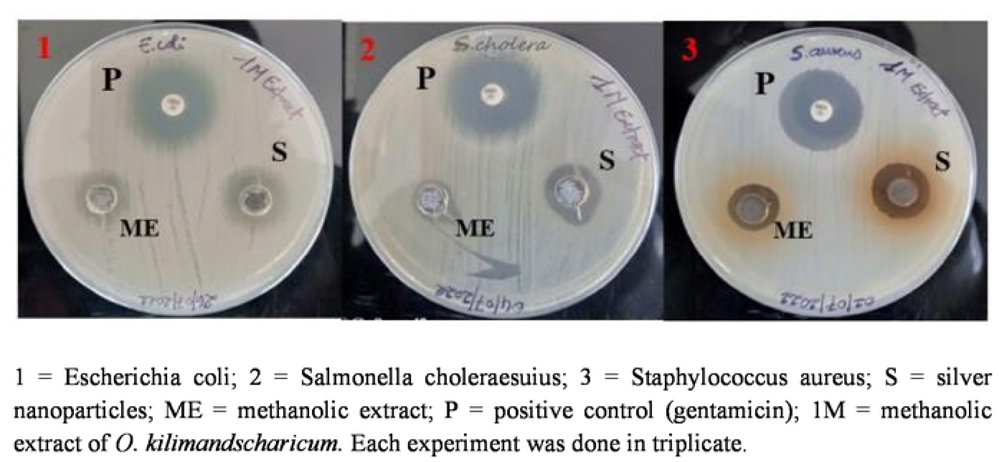
Zones of inhibition for the methanolic extract and silver nanoparticles of the methanolic extract of *O. kilimandscharicum*

**Figure 13.**
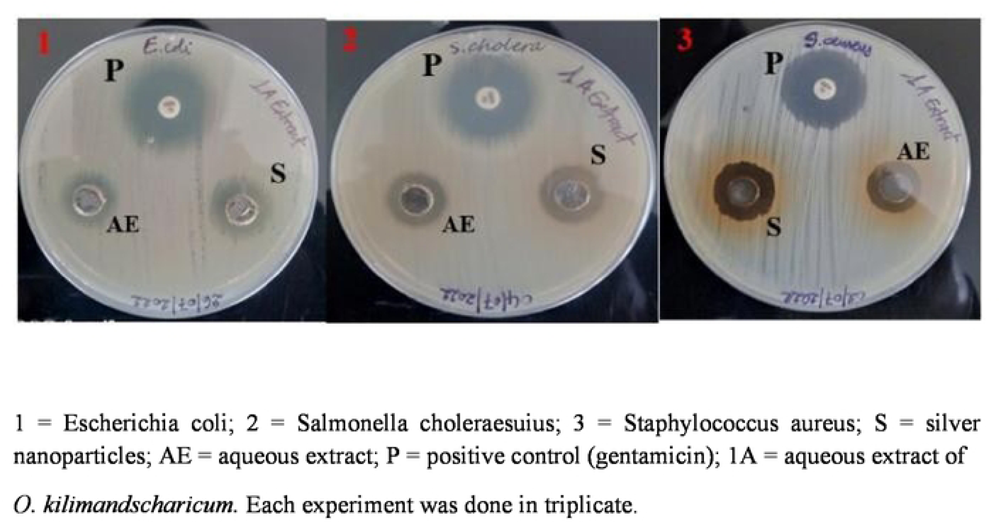
Zones of inhibition for the aqueous extract and silver nanoparticles of the aqueous extract of *O. kilimandscharicum*

**Table 3.**
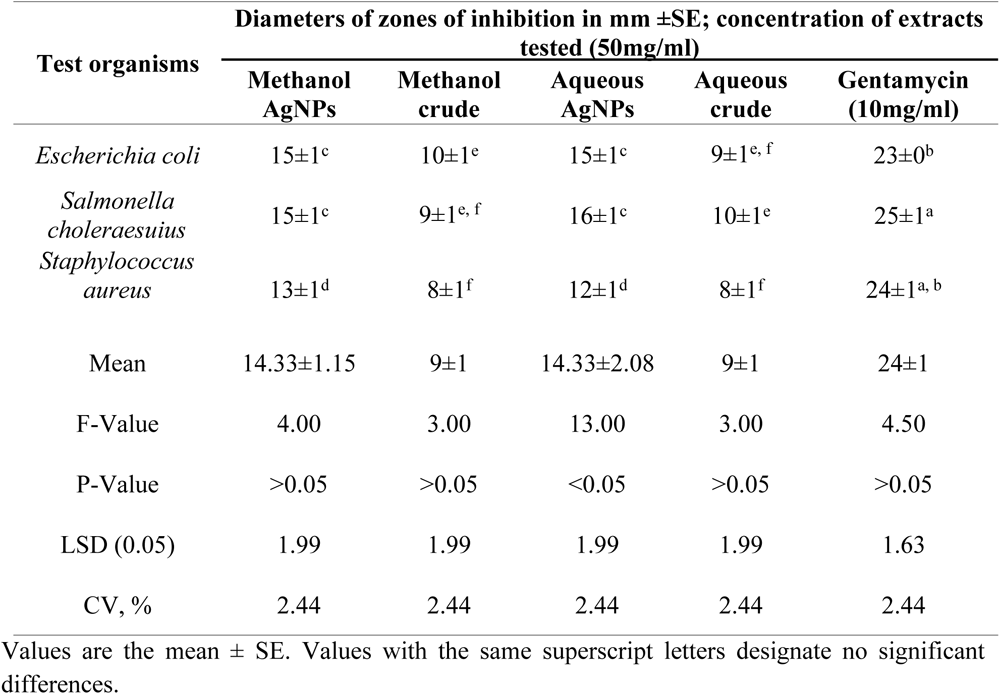
Zone of inhibition against some bacteria strains by methanol extract, aqueous extract and AgNPs of methanol extract, AgNPs of aqueous extract of *Ocimum kilimandscharicum*.

The findings of the zone of inhibition were represented as mm. The data were presented as Mean ± SE of triplicate, as shown in Table 3. Silver nanoparticles made from aqueous extract had the highest zone of inhibition (16±1 mm), whereas water and methanolic extracts had the lowest (8±1 mm). Silver nanoparticles are more effective than crude extracts, as shown in Table 3, where the lowest zone of inhibition of silver nanoparticles is 12±1 mm, and the one for crude extracts is 8±1 mm. Moreover, silver nanoparticles are more effective against Gram-negative bacteria, *Escherichia coli* and *Salmonella choleraesuius* (zone of inhibition between 15 to 16±1 mm). In contrast, they are the least effective on Gram-positive bacteria, *Staphylococcus aureus* (zone of inhibition between 12 at 13±1 mm. ANOVA of aqueous silver nanoparticles (F (2,6) = 13.00, p<0.05) in Table 3 revealed that there was a statistically significant difference between the groups. The superscripted letters in Table 3 indicate significant differences between means.

Several publications describe the processes by which silver nanoparticles function, such as alteration of membrane permeability [27,38], bacterial protein precipitation [50,51], tiny size and vast surface area to attach to the cell wall [41], DNA cleavage [50], production of free radicals, and electrostatic attraction [44,52].

Mariselvam *et al.* [53] and Numan *et al.* [54] consider zones of inhibition with sizes of < 9 mm, 9 - 12 mm, and 13 - 18 mm inactive, moderately active, and active, respectively. This study revealed that silver nanoparticles produced from methanolic and aqueous extracts of *O. kilimandscharicum* and crude extracts have significant antibacterial activity against Gram-positive and negative bacteria (Figure 14 and Table 3). Different zones of inhibition demonstrated that silver nanoparticles were more efficient against Gram-negative bacteria (*S. choleraesuius* and *E. coli*) than against Gram-positive bacteria (*S. aureus*) (Figure 16).

**Figure 14.**
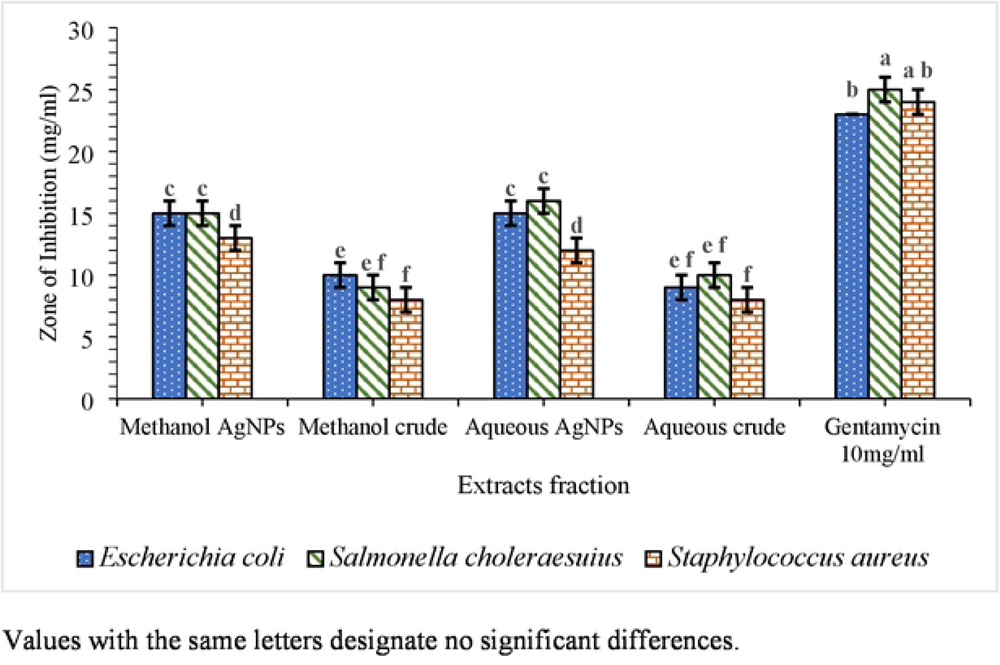
The zone of inhibition for the aqueous and methanolic extracts and their silver nanoparticles of *O. kilimandscharicum* against the indicated bacteria.

Baran *et al.* [44], Khan *et al.* [37], Ahmed *et al.* [55], and Khanal *et al.* [38] all reported identical findings. This may be due to changes in cell wall composition or stiffness between Gram-positive and Gram-negative bacteria [56,57].

In contrast, Phuyal *et al.* [41], Khodadadi *et al.* [58], and Kaur *et al.* [59] demonstrated more antibacterial activity against Gram-positive bacteria (*S. aureus*) than against Gram-negative bacteria (*E.coli* and *S. choleraesuius*).

Khalil *et al*. [60] presented intriguing data about the antibacterial activity of the crude extract of *Ziziphus nummularia* fruit compared to its mediated silver nanoparticles, which aligns with our results. The diameters of the inhibitory zones for the crude aqueous extract of *Ziziphus nummularia* fruit were 11.66 mm for *S. aureus* and 6.30 mm for *E. coli*. For silver nanoparticles, however, it climbed to 16.66 mm for *S. aureus* and 18.66 mm for *E. coli*, respectively [60].

Khanal *et al.* [38] reported root extracts of *Rubus ellipticus* with zone of inhibition values of 10 mm (*E. coli*), 9 mm (*S. aureus*), 10 mm (*K. pneumoniae*), and 11 mm (*E. faecalis*). In contrast, the root extracts of *Rubus ellipticus*-mediated silver nanoparticles were 13 mm (*E. coli*), and 12 (*E. faecalis*). This proved that the antibacterial properties of silver nanoparticles were greater than those of the crude extract [38].

#### 3.2.2. Minimum Inhibitory Concentration (MIC) and Minimum Bactericidal Concentration (MBC)

MIC is the lowest concentration at which no colour change occurs [61,62], whereas MBC is the lowest concentration at which no growth is seen on a petri dish [63].

A, C & E = silver nanoparticles; B, D & F = methanolic extract; 1-12 = wells; P = positive control (bacteria without extract); N = negative control (Mueller Hinton Broth without bacteria)

The findings of the MIC and MBC were represented as mg/ml. The data were presented as Mean ± SE of triplicate, as shown in Table 4. Figures 15, 16, 17a, 17b, 18a, 18b, 19a, 19b, 20 and Table 4 show the findings, respectively. Silver nanoparticles made from aqueous and methanolic extracts had the lowest MIC (6.25±0.00 mg/ml), whereas water and methanolic extracts had the highest MIC (25±0.00 mg/ml). Silver nanoparticles made from aqueous and methanolic extracts had the lowest MBC (12.5±0.00 and 25±0.00 mg/ml), whereas water and methanolic extracts had the highest MBC (50±0.00 mg/ml). Silver nanoparticles are more effective than crude extracts, as shown in Table 4, where the MIC of silver nanoparticles is 6.25±0.00 mg/ml, and the one for crude extract is 25±0.00 mg/ml. As we said above, silver nanoparticles are more effective on Gram-negative bacteria*, Escherichia coli* and *Salmonella choleraesuius* (MBC 12.5±0.00 mg/ml). In contrast, they are the least effective on Gram-positive bacteria, *Staphylococcus aureus* (MBC, 25±0.00 mg/ml). Only ANOVA of MBCs’ aqueous and methanol silver nanoparticles (F (2,6) = Infinity, p<0.0001) in Table 4 revealed that there was a statistically significant difference between the groups. The superscripted letters in Table 4 indicate significant differences between means.

**Figure 15.**
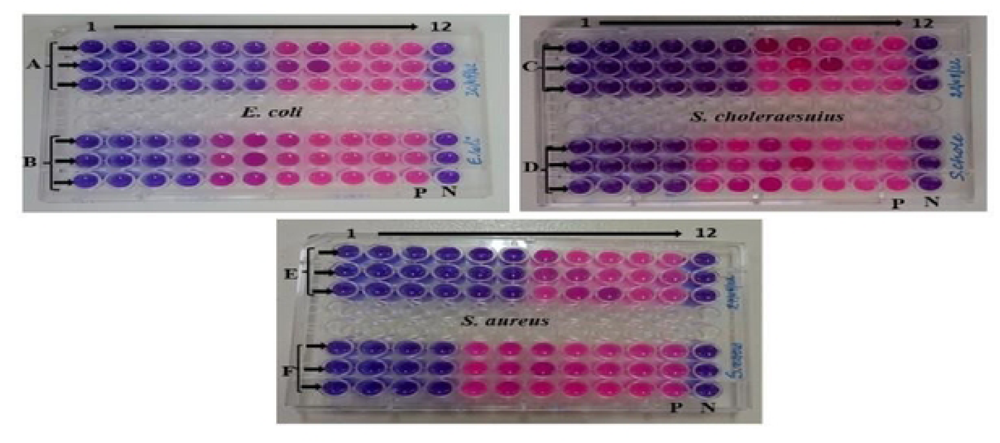
Minimum inhibitory concentrations of the methanolic extract and silver nanoparticles of *O. kilimandscharicum* against the indicated bacteria.

**Figure 16.**
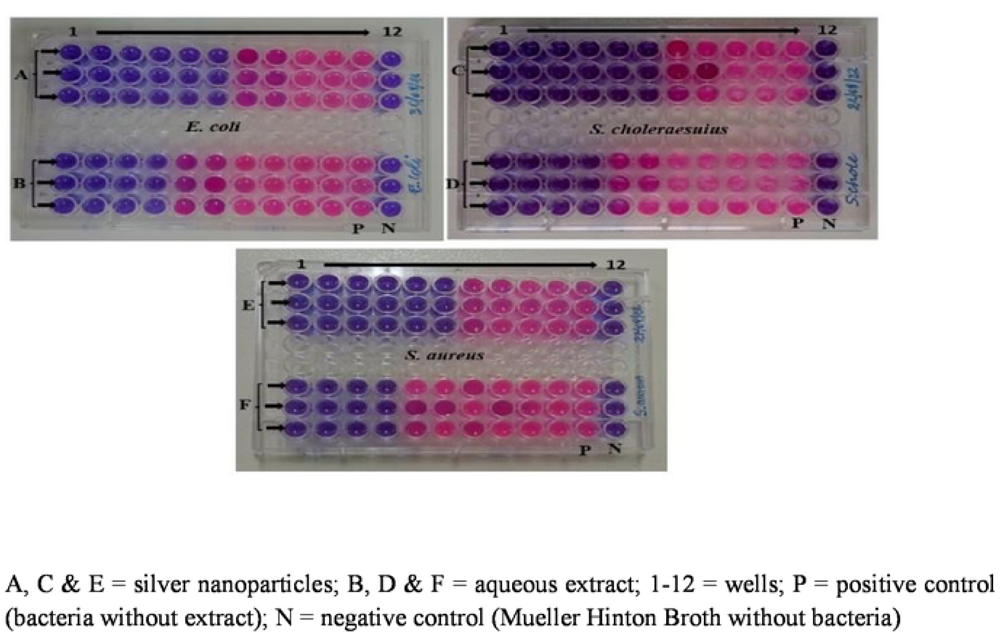
Minimum inhibitory concentrations of the aqueous extract and silver nanoparticles of *O. kilimandscharicum* against the indicated bacteria.

**Figure 17a.**
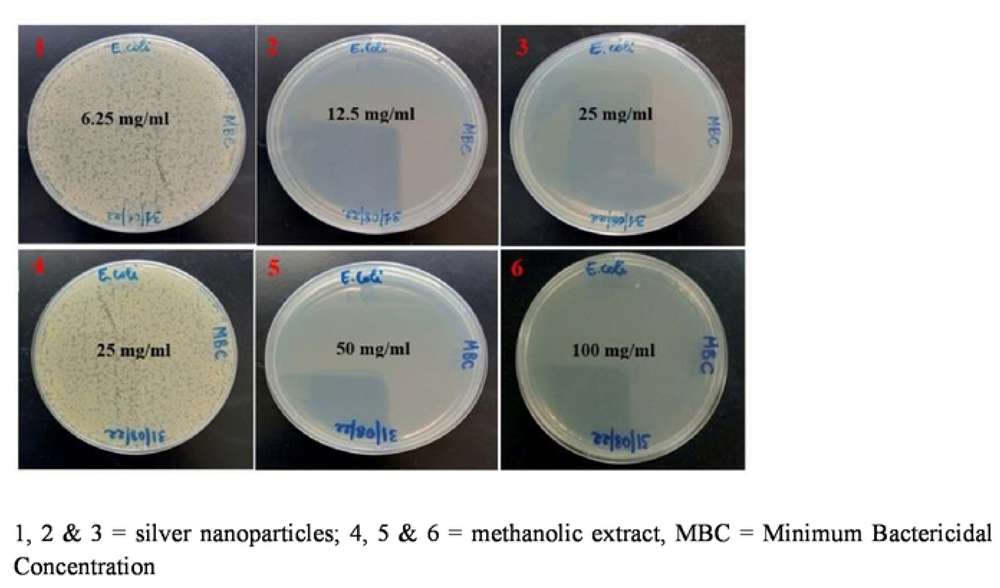
Minimum bactericidal concentrations of the methanolic extract and silver nanoparticles of *O. kilimandscharicum* against *E. coli*

**Figure 17b.**
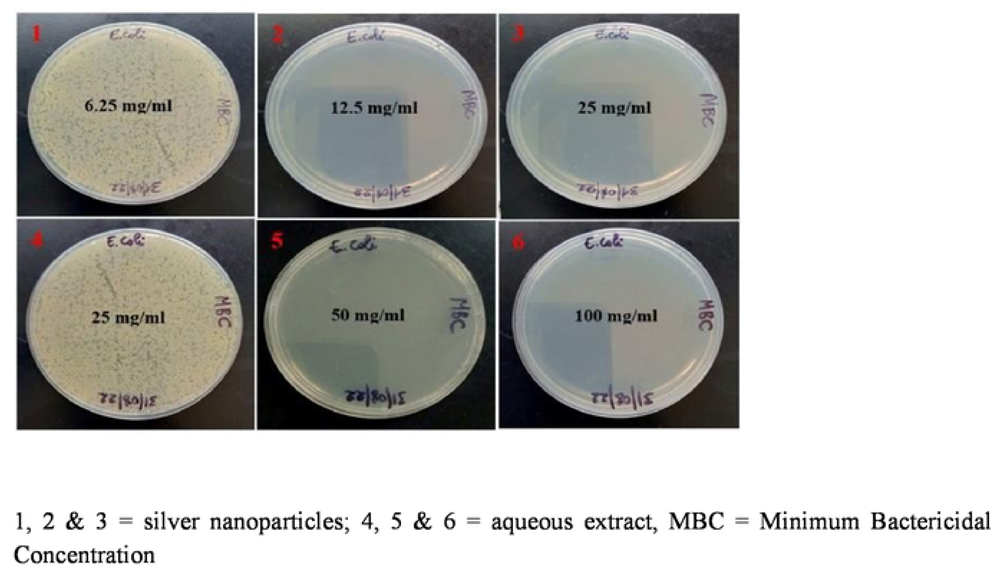
Minimum bactericidal concentrations of the aqueous extract and silver nanoparticles of *O. kilimandscharicum* against *E. coli*

**Figure 18a.**
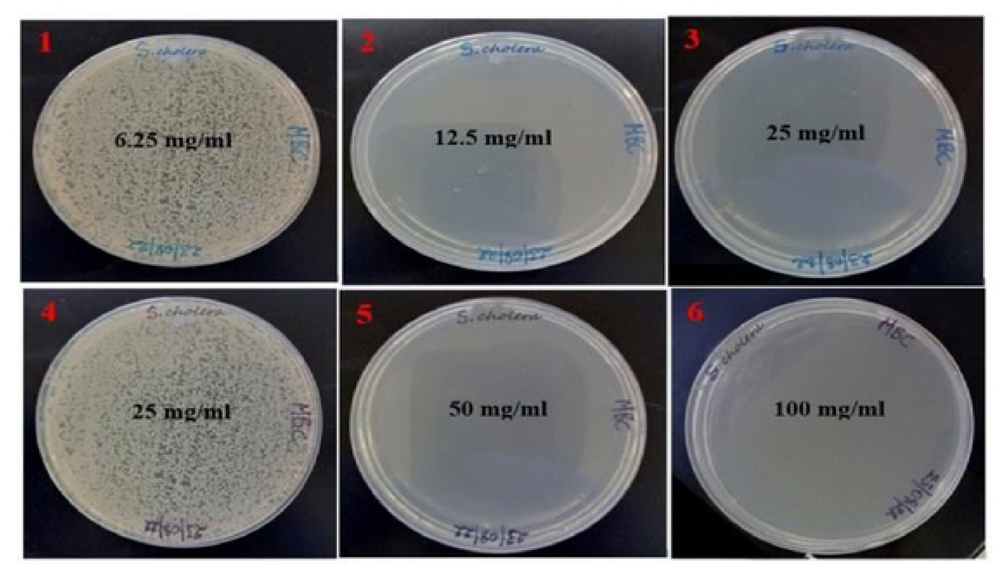
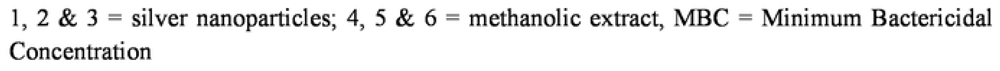
Minimum bactericidal concentrations of the methanolic extract and silver nanoparticles of *O. kilimandscharicum* against *Salmonella choleraesuius*

**Figure 18b.**
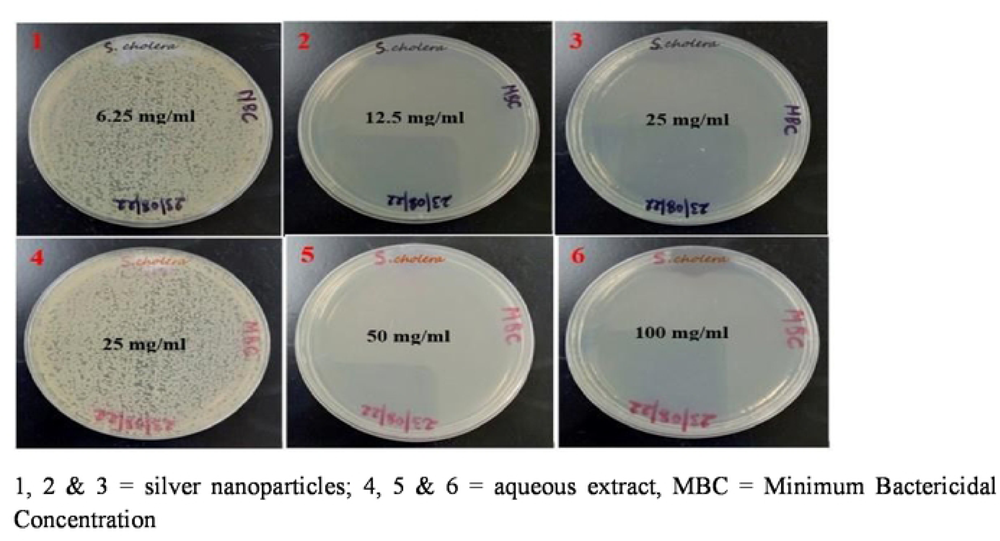
Minimum bactericidal concentrations of the aqueous extract and silver nanoparticles of *O. kilimandscharicum* against *Salmonella choleraesuius*

**Figure 19a.**
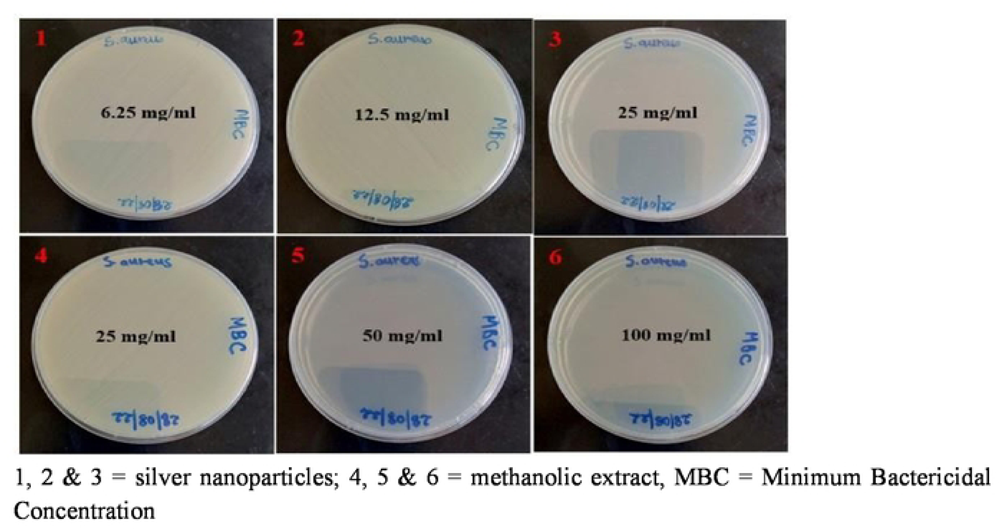
Minimum bactericidal concentrations of the methanolic extract and silver nanoparticles of *O. kilimandscharicum* against *Staphylococcus aureus*

**Figure 19b.**
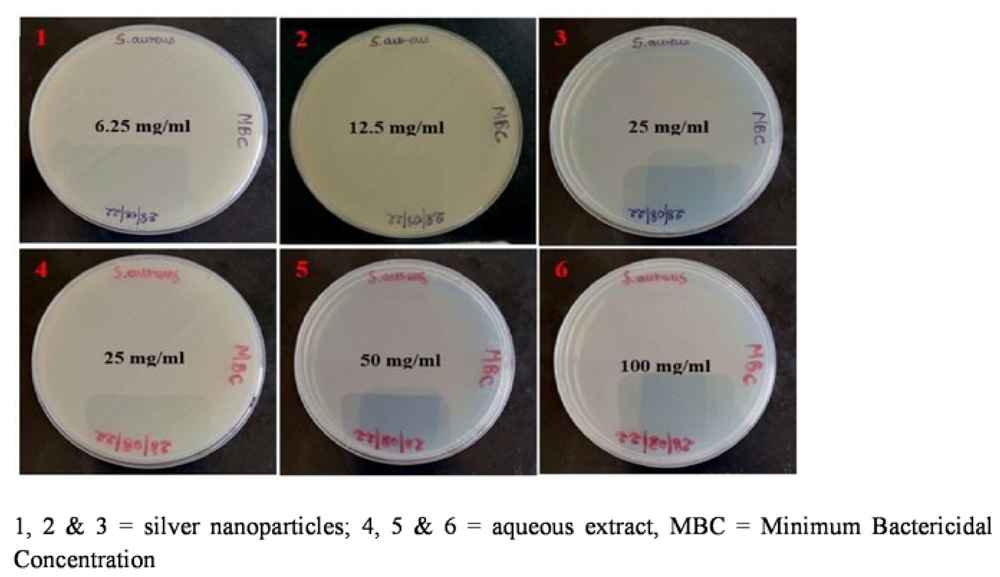
Minimum bactericidal concentrations of the aqueous extract and silver nanoparticles of *O. kilimandscharicum* against *Staphylococcus aureus*

**Figure 20.**
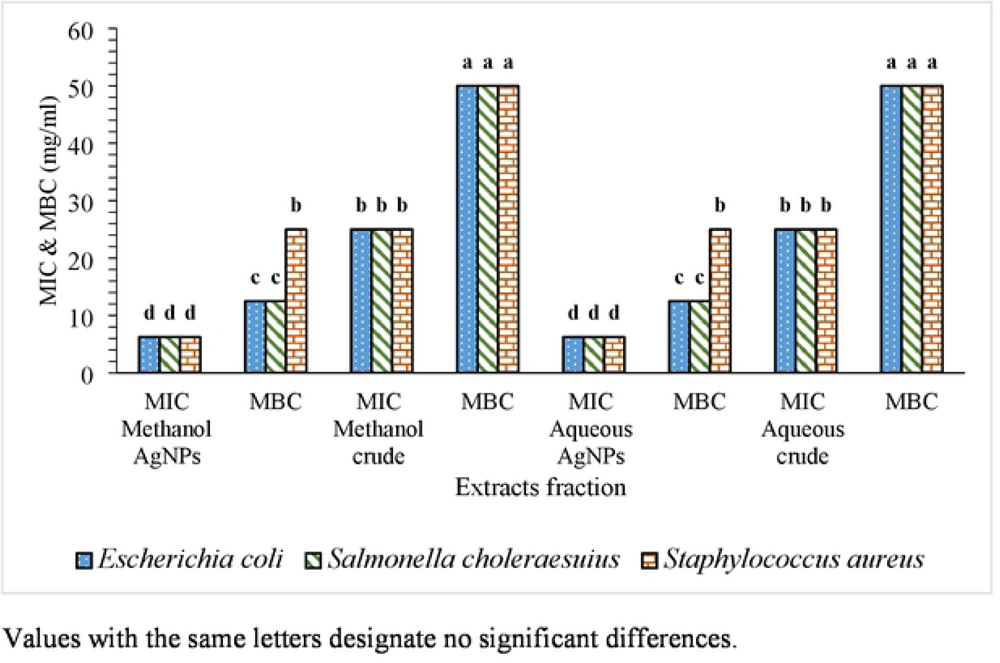
The MIC and MBC for the aqueous and methanolic extract and their silver nanoparticles of *O. kilimandscharicum* against the indicated bacteria.

**Table 4.** MIC (mg/ml) and MBC (mg/ml) ±SE against some bacteria strains by methanol extract, aqueous extract and AgNPs of methanol extract, AgNPs of aqueous extract of Ocimum kilimandscharicum.

| Test organisms | Methanol AgNPs |  | Methanol crude |  | Aqueous AgNPs |  | Aqueous crude |  |
| --- | --- | --- | --- | --- | --- | --- | --- | --- |
|  | MIC | MBC | MIC | MBC | MIC | MBC | MIC | MBC |
| *Escherichia coli* | 6.25±0.0<sup>d</sup> | 12.5±0.0<sup>c</sup> | 25±0.0<sup>b</sup> | 50±0.0<sup>a</sup> | 6.25±0.0<sup>d</sup> | 12.5±0.0<sup>c</sup> | 25±0.0<sup>b</sup> | 50±0.0<sup>a</sup> |
| *Salmonella choleraesuius* | 6.25±0.0<sup>d</sup> | 12.5±0.0<sup>c</sup> | 25±0.0<sup>b</sup> | 50±0.0<sup>a</sup> | 6.25±0.0<sup>d</sup> | 12.5±0.0<sup>c</sup> | 25±0.0<sup>b</sup> | 50±0.0<sup>a</sup> |
| *Staphylococcus aureus* | 6.25±0.0<sup>d</sup> | 25±0.0<sup>b</sup> | 25±0.0<sup>b</sup> | 50±0.0<sup>a</sup> | 6.25±0.0<sup>d</sup> | 25±0.0<sup>b</sup> | 25±0.0<sup>b</sup> | 50±0.0<sup>a</sup> |
| Mean | 6.25±0.00 | 16.66±7.21 | 25±0.0 | 50±0.0 | 6.25±0.0 | 16.66±7.21 | 25±0.0 | 50±0.0 |
| F-Value | - | Infinity | - | - | - | Infinity | - | - |
| P-Value | - | <0.0001 | - | - | - | <0.0001 | - | - |
| LSD (0.05) | 0 | 0 | 0 | 0 | 0 | 0 | 0 | 0 |
| CV, % | 2.44 | 2.44 | 2.44 | 2.44 | 2.44 | 2.44 | 2.44 | 2.44 |

Values are the mean ± SE. Values with the same superscript letters designate no significant differences

Abdellatif *et al.* [64] showed a similar scenario employing an aqueous extract of *Thymus vulgaris* leaves, *Zingiber officinale* roots, and their silver nanoparticles against the *S. aureus* clinical strain. The MBC value of aqueous *Thymus vulgaris* leaf extract was 0.2825 mg/ml, which decreased to 0.0706 mg/ml in silver nanoparticles. For aqueous preparations of *Zingiber officinale* root extract, the concentration decreased from 2,2600 mg/ml to 0.5650 mg/ml [64].

Several research investigations, such as Periasamy *et al.* [65], Periyasami *et al.* [66], Phuyal *et al.* [41], and Khanal *et al.* [38], have shown the occurrence of silver nanoparticles with strong biological activity.

## 4. Conclusion

This work describes the manufacture of silver nanoparticles from aqueous and methanolic O. kilimandscharicum leaf extract. UV–Vis revealed silver nanoparticles’ surface plasmon resonance. Biosynthesized AgNPs demonstrated significant antibacterial action against all pathogens. To combat rising concerns, AgNPs will make a considerable contribution to antimicrobial agent research. We suggest that the work can be continued to examine the *in vitro* and *in vivo* toxicity of silver nanoparticles generated sustainably.

## Conflict of interest

There are no stated conflicts of interest by the authors.

## Acknowledgements

The research from which this dataset was obtained was funded by the Pan African University Institute for Basic Sciences, Technology and Innovation Doctoral grant to HSO, MB401-0005/19. The Pan African University of Basic Science, Technology, and Innovation provided the facilities and laboratory support necessary for the execution of this experiment, which the authors gratefully recognize. They are quite appreciative of the African Union’s assistance with research money. The bacteria strains were generously provided by the Kenya Medical Research Institute (KEMRI), for which the authors are very grateful.

## Notes

### Competing Interest Statement

The authors have declared no competing interest.

